# Movement Decoding using Spatio-Spectral Features of Cortical and Subcortical Local Field Potentials

**DOI:** 10.1101/2021.06.06.447145

**Authors:** Victoria Peterson, Timon Merk, Alan Bush, Vadim Nikulin, Andrea A Kühn, Wolf-Julian Neumann, Mark Richardson

## Abstract

The application of machine learning to intracranial signal analysis has the potential to revolutionize deep brain stimulation (DBS) by personalizing therapy to dynamic brain states, specific to symptoms and behaviors. Most decoding pipelines for movement decoding in the context of adaptive DBS are based on single channel frequency domain features, neglecting spatial information available in multichannel recordings. Such features are extracted either from DBS lead recordings in the subcortical target and/or from electrocorticography (ECoG). To optimize the simultaneous use of both types of signals, we developed a supervised online-compatible movement decoding pipeline based on multichannel and multiple site recordings. We found that adding spatial information to the model has the potential to improve decoding. In addition, we demonstrate movement decoding from spatio-spectral features derived from cortical and subcortical oscillations. We demonstrate between-patients variability of the spatial neural maps and its relationship to feature decoding performance. This application of spatial filters to decode movement from combined cortical and subcortical recordings is an important step in developing machine learning approaches for intelligent DBS systems.

## 1 Introduction

Movement decoding from invasive neural oscillations has the potential to change the way implantable braincomputer interfaces (BCI) can aid the therapy of movement disorders. Deep brain stimulation (DBS) of the subthalamic nucleus (STN) or the globus pallidus internus (GPi) is an effective alternative treatment for patients with Parkinson’s disease (PD) [1,2]. Although continuous high-frequency DBS consistently improves motor symptoms of PD patients [3], several side effects, such as dysarthria, dyskinesia and balance issues [4,5] can complicate treatment. In order to minimize such side-effects, closed-loop or adaptive DBS (aDBS), guided by the patient’s electro-clinical state, has been proposed as a strategy to reduce the amount of unnecessary stimulation energy delivered to the brain [6]. Closed-loop DBS devices therefore are bidirectional invasive brain-computer interfaces (BCI) that can adapt stimulation in dependence of control algorithms that are informed by brain signals.

The viability and success of bidirectional BCIs for aDBS strongly depends on the identification of reliable biomarkers reflecting patients’ symptom severity and change with treatment, as well as on the computational strategies used for neural decoding of such states and behavior. This will allow to augment aDBS strategies with machine learning, e.g. by decoding kinematic parameters which could be used in the future to refine stimulation parameters [7]. It has been shown that behavioral neural biomarkers can be identified from local field potentials (LFPs) in the STN [8,9], electrocorticography (ECoG) [10] or the combinational use of both [11,12].

Making use of the spatio-spectral information has become an established approach in the field of non-invasive BCI research, where the spatial information of the sensors can be incorporated into the decoding algorithms. Importantly, due to the overlapping activity of multiple sources, specific neural population activity cannot be observed directly and must be inferred with uncertainty [13]. Statistical generative models often assume that brain signals arise from activity of uncorrelated sources, and that such sources appear distorted in the signal recording as a consequence of a linear mixing (due to volume conduction), as illustrated in Figure 1. While the spatial signal to source relationship is more precise in the field of invasive neurophysiology, the same underlying mechanisms are at play, and invasive recordings are contaminated by a mix of local and distant (volume conducted) brain activity.

**Figure 1:**
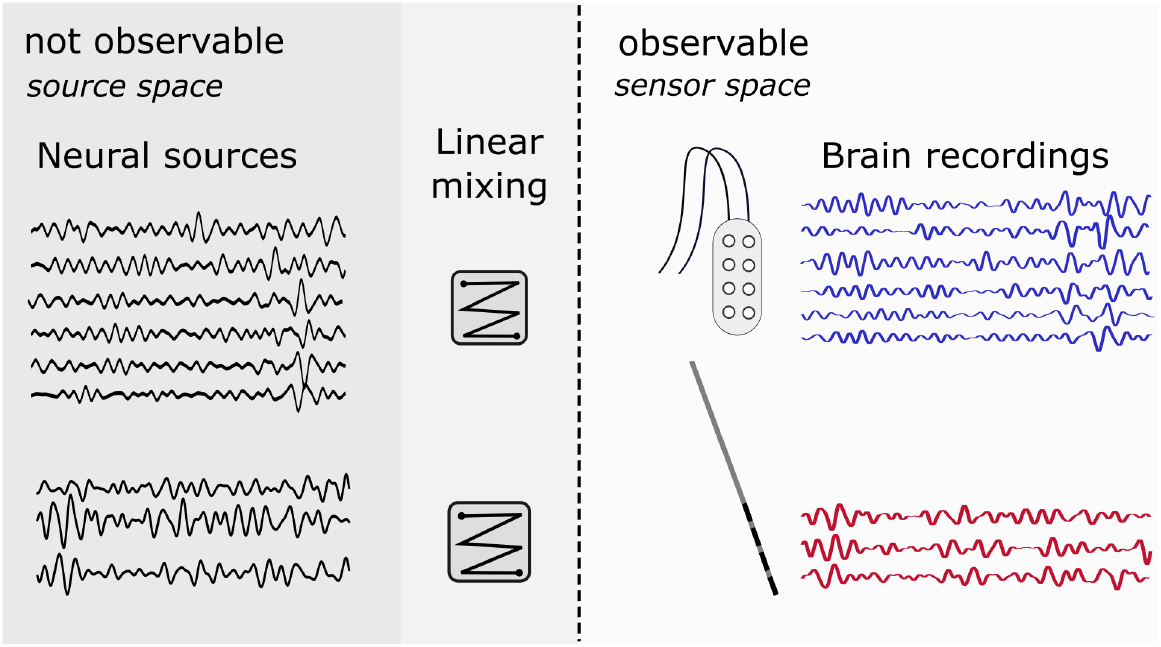
Illustration of the generative statistical model in invasive neurophysiology. The true neural sources are inferred by unobservable source space signals. These sources are mixed to constitute the sensor space signals (e.g. ECoG/STN-LFP). Spatial filters aim at estimating the sources by projecting the data in the sensor space to the source space.

Spatial filtering methods aim to extract the relevant spatial information embedded in multivariate (multichannel) signals. They directly work in the channel-time space of the signal, capturing spatial relationships of the brain recordings. In the particular case of statistical spatial filters (Figure 1), the unobservable neural sources are estimated via decomposition of the multichannel brain recording, i.e., by demixing the observable brain signals. The source power comodulation (SPoC) method [14] is a supervised spatial filter approach which allows extraction of band-power related features correlated to a target signal (i.e. kinematic parameters, reaction times, etc). Therefore, SPoC is well suited to extract spatio-spectral features for identifying neural biomarkers related to movement, as recently showed in [15].

In this study, we constructed subject-specific ML-based invasive neurophysiology decoding models based on both the frequency and the spatial dimensions of the recorded brain activity, using a strategy that combines a filter-bank analysis with a spatial filtering approach. A multiple recording site (STN-LFP and ECoG) dataset from eleven (11) PD patients who performed a hand movement task during awake DBS implantation surgery was used. The task comprised both contra- and ipsilateral movements with respect to the electrode localization. A generalized linear model (GLM) with Poisson-like regularized regression was implemented for predicting the movement. We evaluated regression performance of the model in two modalities: i) single recording site, in which either ECoG or STN-LFP recordings were used as inputs to the decoding model and ii) multiple recording sites, in which both ECoG and STN-LFP recordings were used together to train the model. Since spatial filters were used, the solution was subjected to neurophysiological interpretation. We analyzed how band-power features and brain recording modalities contribute to movement decoding. We found that multichannel approaches have the potential to improve movement decoding as compared to a single-channel approach. This study advances the use of ML methods with multichannel and multiple site recordings for the potential development of intelligent aDBS devices, taking into account the spatial information of recording sites. The source code used throughout this work is available at GitHub https://github.com/Brain-Modulation-Lab/Paper_SpatialPatternsMovementDecoding.

## 2 Results

### 2.1 Spatio-spectral decomposition for movement decoding in PD patients

We created a spatio-spectral pipeline for movement decoding in the context of invasive neuromodulation. The pipeline is designed to work in real-time scenarios. Thus, data packets (segments) of 1000 ms were used to decode the incoming target sample. These recording segments, either from ECoG, STN-LFP, or the combination of both, were decomposed into 8 well-defined frequency-bands, ranging from theta to gamma band. The SPoC method was used to estimate the neural source most correlated to the hand movement target signal at each frequency band. Then a band-power feature was extracted over the estimated source. Thus, via SPoC, one spatio-spectral feature per frequency-band was extracted. Concatenated features were then used to feed a GLM with Poisson sparse regularized regression to predict the target. See Figure 2 and the *Spatio-spectral multiple recording site decoding pipeline* in Materials and Methods for a detailed description of the model.

**Figure 2:**
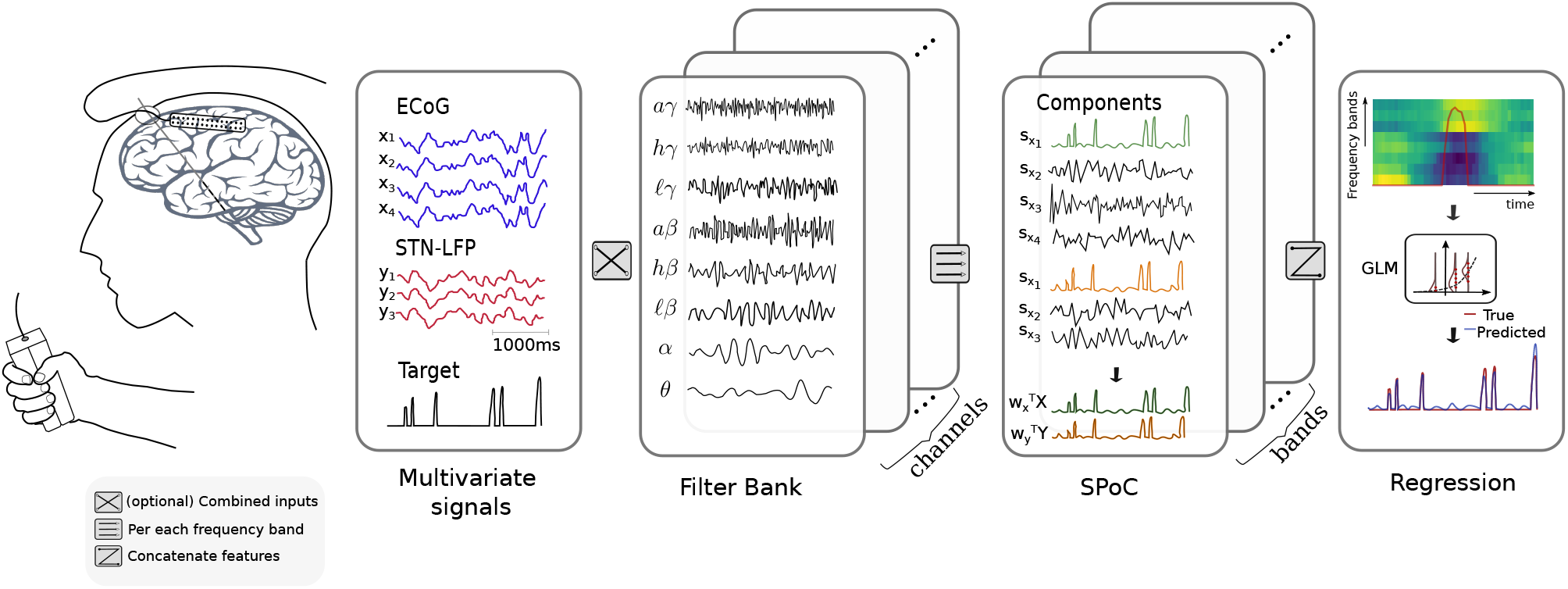
Schematic representation of the decoding pipeline. Segments of 1000 ms from the invasive multichannel recording were decomposed in eight frequency bands. The supervised spatial filtering SPoC method was applied at each frequency band, extracting the source more correlated to the hand grip movement (target variable). One spatio-spectral feature was extracted from each frequency band. Movement decoding was based on a GLM regression model, in which a Poisson sparse penalized regularization was implemented.

### 2.2 Spatio-spectral features inform movement in PD patients

The SPoC method is designed to decompose, in a supervised manner, the multivariate brain recordings into a set of source components. In theory, if enough training data is available for building the decoding model, performance should be superior to that of a univariate approach. The advantages of using spatial approaches are well-known for surface EEG based decoding [14], but given the more immediate relationship of sensor and source, the impact of spatial methods for invasive neurophysiology remains to be elucidated. Here, we compared the decoding capacity of our proposed spatio-spectral decoding model to a single channel pipeline [16]. While the former uses all channels at once to build the model, the latter needs to evaluate the decoding capacity of each electrode individually and then choose the best electrode to run the movement prediction. The comparative results are shown in Figure 3.

**Figure 3:**
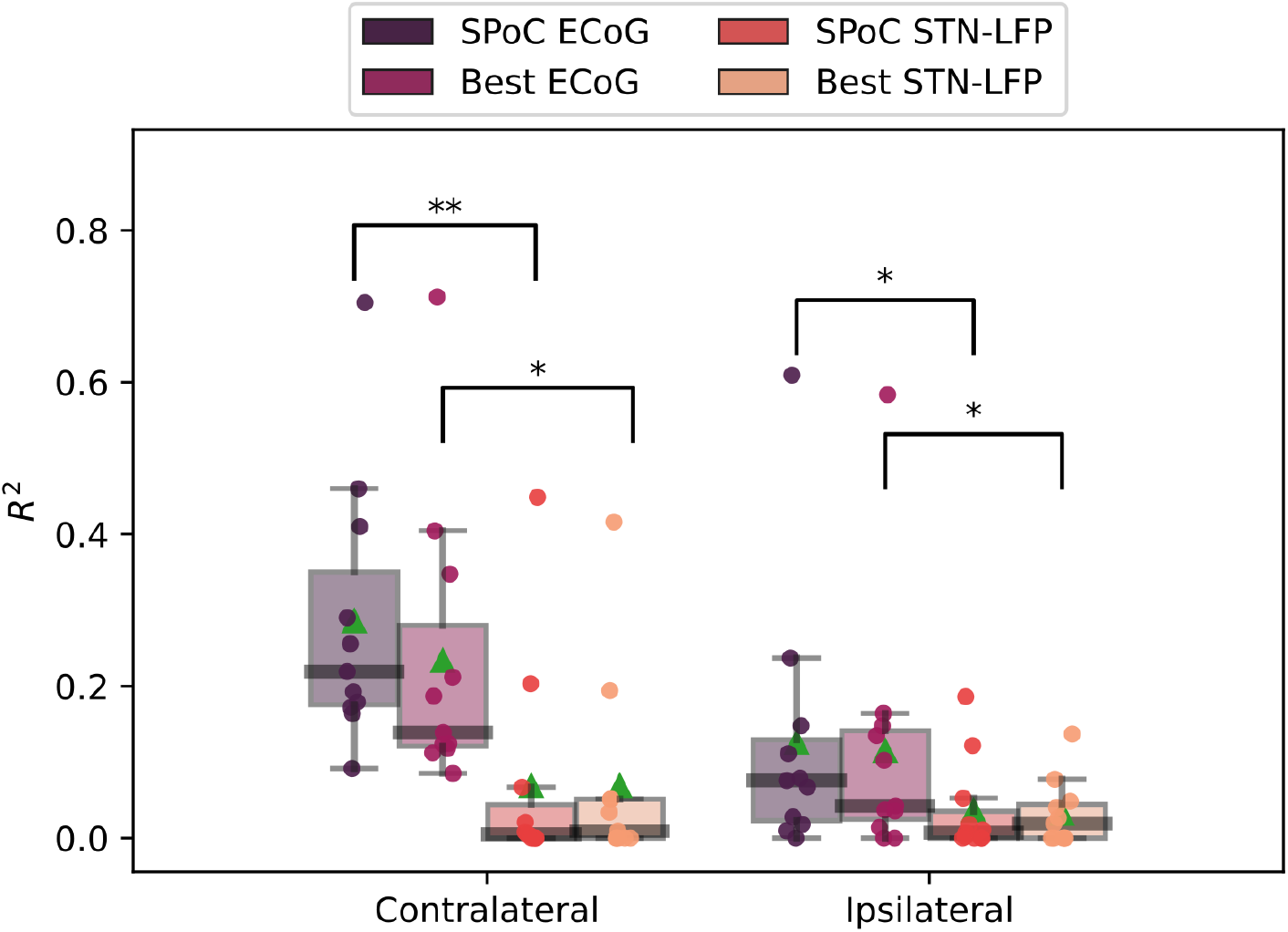
Spatio-spectral brain signal decoding approaches can be applied in invasive neurophysiology. Performance comparison between the spatio-spectral approach (based on SPoC) and the single-channel (Best single recording location) approach, evaluated for each type of brain signal: ECoG and STN-LFP. The performance in decoding contralateral and ipsilateral movement is shown separately. Statistical significance according to Fried-man+Nemenyi test is denoted by ** (p-value ≤ 0.01), * (0.01 < p-value ≤ 0.05). The green triangle indicates the mean value.

We first found that a decoding model based on contralateral movements performed better than its ipsilateral counterpart (e.g., mean *R*^2^ SPoC ECoG contralateral: 0.29, mean *R*^2^ SPoC ECoG ipsilateral: 0.13, p-value¡0.01). Second, decoding based only on ECoG recording was consistently better than that based only on STN-LFP signals (e.g., mean *R*^2^ SPoC ECoG contralateral: 0.29, mean *R*^2^ SPoC STN-LFP contralateral: 0.07, p-value¡0.01). Third, the spatial information seemed to have the potential to improve the decoding capacity of the model (e.g., mean *R*^2^ SPoC ECoG contralateral: 0.29, mean *R*^2^ Best ECoG contralateral: 0.23, p-value=0.056).

### 2.3 Combining cortical and subcortical spatio-spectral features does not improve movement decoding

ECoG and STN-LFP signals were recorded simultaneously in these subjects, allowing the definition of decoding models trained on a combination of both recording site signals. The spatio-spectral features were extracted from 8 frequency bands, resulting in overall 16 input features (8 features per modality). Figure 4 shows the performance of the decoding model using ECoG + STN-LFP recordings against decoding models that only use either ECoG or STN data. The statistical analysis indicates that there are no significant differences between the combined ECoG+ STN-LFP decoding model and the decoding model based only on ECoG signals (p-value=0.9).

**Figure 4:**
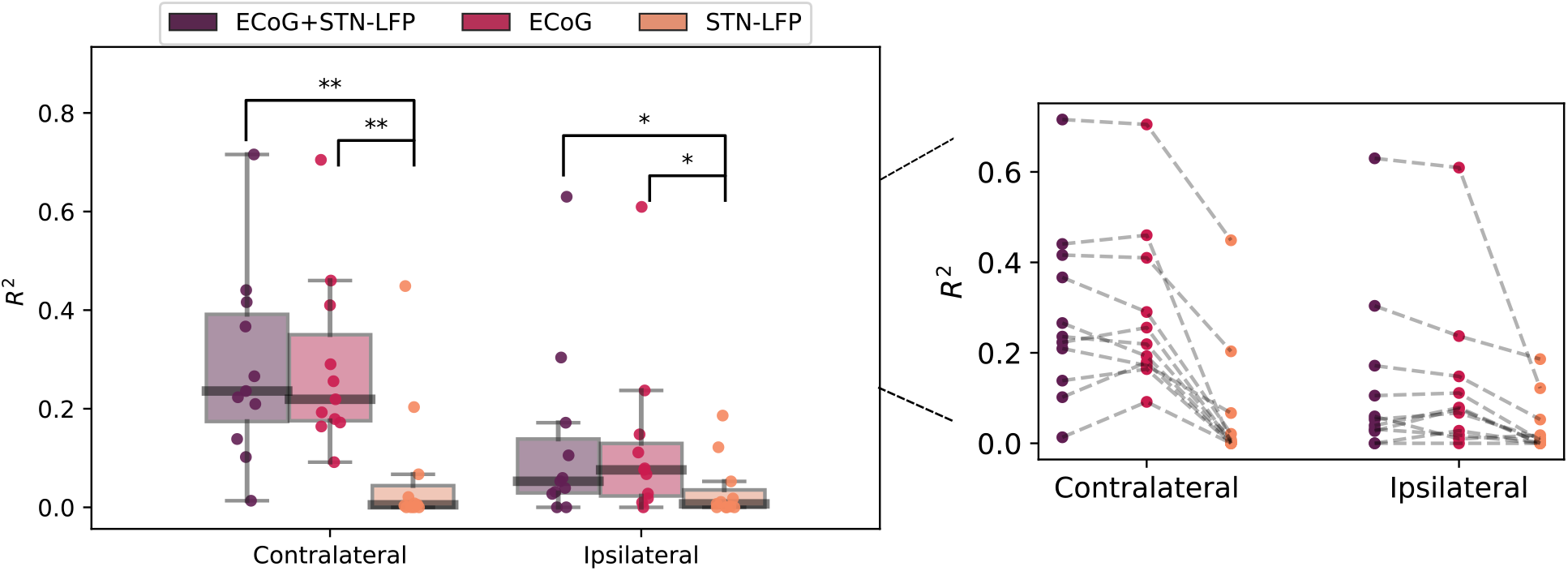
Adding STN-LFPs features does not significantly improve ECoG-based decoding. Performance comparison between multiple recording sites and single recording site approaches. The performance in decoding contralateral and ipsilateral movements is shown separately. The plot on the right links decoding performance across the same subject (dot) when using STN, ECoG and ECoG + STN-LFP data. Statistical significance according to Friedman+Nemenyi test is denoted by ** (p-value ≤ 0.01), * (0.01 < p-value ≤ 0.05).

#### 2.3.1 Model interpretability of cases in which the combined ECoG+STN models outperformed models trained on ECoG signals only

Although on average combining spatio-spectral features from cortex and subcortex recording did not outperform the approach using only ECoG, we further analyzed those cases in which the combinational use did improve movement decoding. Thus, for three patients in which performance improvement was found by adding the STN signal to the ECoG decoding model, a detailed model interpretability analysis was performed.

Figure 5 shows the solutions vector of the Poisson-like GLM together with the learned spatial patterns from the cortex and subcortex signals at each frequency band considered. Interestingly, for these subjects, the decoding model depends on spatio-spectral features coming from both cortical and subcortical regions. The spatial patterns from selected features map movement-related brain activity.

**Figure 5:**
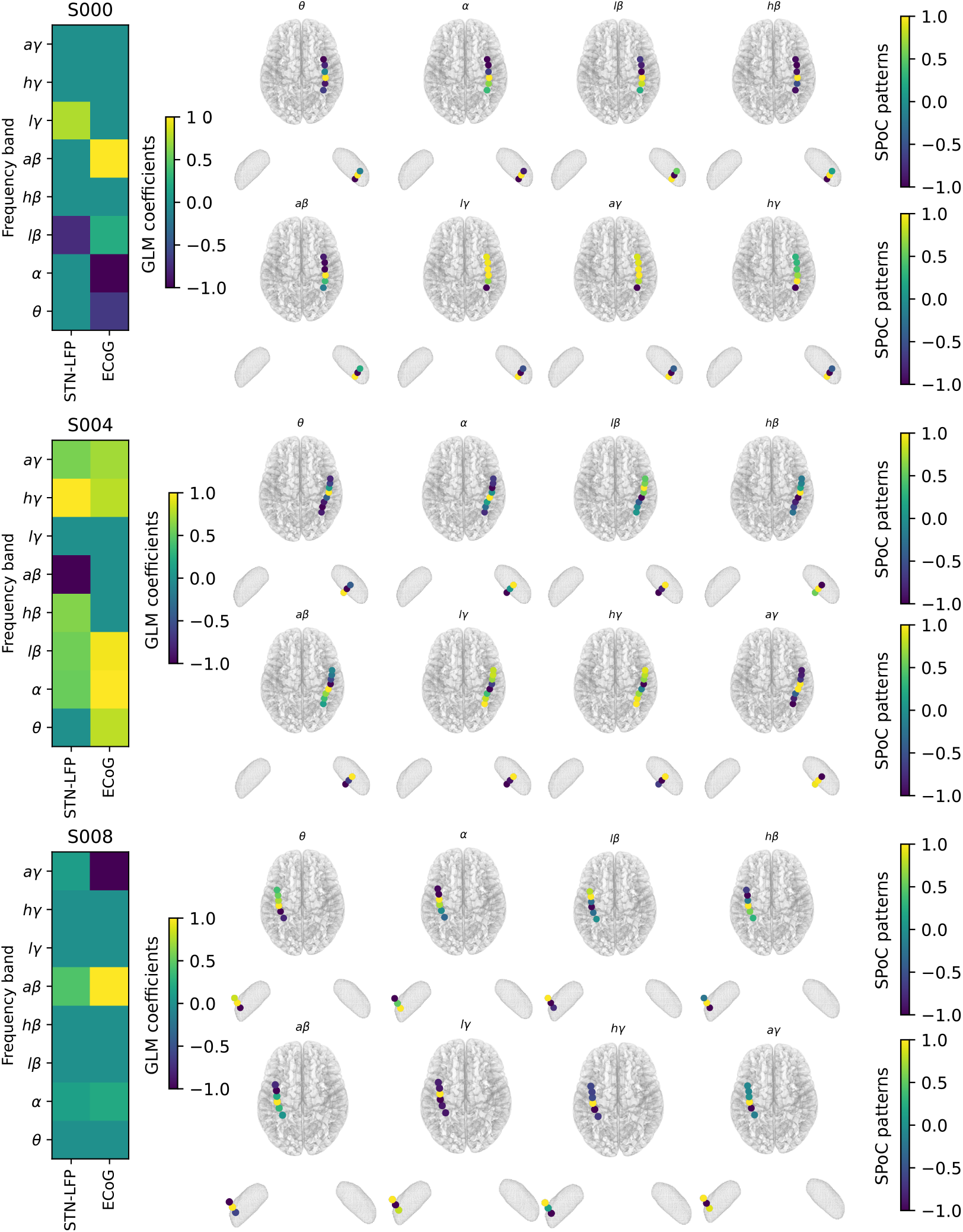
Spatio-spectral information shows feature importance and maps brain activity related to gripforce encoding. Model interpretability for three subjects (S000, S004, S008) in which the multisite recording site outperforms the single site ECoG decoding capacity. One subject per figure subpanel. Left side: GLM coefficient values; right side: the associated spatial pattern at each frequency band considered. Due to the nature of the Poisson-like distribution, for better interpretation of the output, the coefficient values were transformed by taking the exponential and then normalized. For illustration purposes, the spatial patterns are also normalized between -1 and 1.

## 3 Discussion

In this study we implemented a spatio-spectral approach to decode movement using invasive brain recordings in patients with Parkinson’s disease. The supervised spatial filter SPoC method was used to extract features at eight pre-defined frequency bands. The predictive model was built via GLM with regularized Poisson-like sparse regression. Contra- and ipsilateral movements were decoded separately. First, the spatio-spectral approach was compared against a single channel approach. Second, the decoding pipeline was tested using combined ECoG and subthalamic LFP recordings and compared to models only informed by ECoG. Model interpretability was assessed by analyzing the cases in which improvement was found at the combined recording site approach.

### 3.1 Spatial information in invasive neurophysiology

The use of blind source separation methods has been largely investigated for understanding brain networks by using different brain imaging modalities. These methods have one objective in common: recovering the unknown (unobservable) source signals that explain the observable brain activity [17]. The statistical generative models on which the spatial filtering methods rely, assume that the recorded brain activity is a consequence of the distributions of different rhythms across electrode contacts (see Figure 1).

In invasive neurophysiology, as compared to non-invasive electro- or magnetoencephalography, the recordings better reflect the surface distribution of cortical sources, yet there is still a distortion effect between the activity of sources and the recordings [18]. In this work, we used a supervised spatial filtering approach for finding the neural source more correlated in power to the movement task. The performance of this spatial filtering method was compared against a similar decoding model which relies on correlating the power of a single recording location with movement [16]. Our results showed that introducing the spatial information to a band-power decoding model has the potential to improve decoding capacity in invasive neurophysiology (see Figure 3). In particular, this improvement was observed for the ECoG recording but not for the STN-LFP. This could be explained by the fact that in our cohort the STN-LFP were recorded by using 4 contacts separated by 0.5 mm (see *Data* in Materials and Method), and thus minimal relative spatial information could be extracted. Similarly, most of the ECoG recordings were acquired by using low-dimensional ECoG grids (6 or 8 channels). We argue that higher improvement could potentially be found for larger high-density ECoG electrodes.

We would like also to note here the superiority of ECoG over STN-LFP for movement decoding (see figures 3 and 4). These findings are in line with Merk et al. [16] and could be explained by the fact that ECoG recordings have a better spatial resolution (more electrode contacts), providing a much stronger signal and allowing measurements of higher amplitude oscillations than depth recordings [19].

### 3.2 Combining spatio-spectral features from multiple site recordings

Several works have investigated the relationship between STN oscillations and those from cerebral cortex in patients with PD. It has been shown that the beta band of the STN is coupled with the cortex activity in PD patients, and that such coupling changes with medication level [20]. Recently, the phase-locking between STN oscillations and cortical beta oscillations has been reported to be different with respect to the cortical areas being studied [21]. Thus, we investigated how the combinational use of spatio-spectral features coming from simultaneously recorded cortical and subcortical activity could impact decoding performance. On average, we found no gain when adding the STN-LFP features to the ECoG approach with respect to using only ECoG features. The fact that no improvement was found in the combinational use of STN-LFP and ECoG features, however, could reflect the fact that we explored here only features that rely on the band power information of the brain oscillations. In fact, recently coherence between STN and motor cortex was shown to be relevant for differentiating motor states in PD patients [12].

Interestingly, there were cases in which the combination of ECoG and LFP recordings approach did improve the decoding capacity when compared to models informed by ECoG alone. To understand the underlying mechanisms that led to such improvement, the decoding model for those particular subjects was analyzed. One of the main advantages of using blind-source separation methods for finding the neural sources is that the solution is neurophysiologically interpretable. Since a linear model was used to learn the mapping between features and movement target output, the coefficient values of the solution can depict feature importance. Figure 5 shows, for each subject, the solution coefficient values associated to each spatio-spectral feature across the different brain modalities and frequency bands. Common to these subjects is that both ECoG and STN coefficients show a strong spatial pattern, with a frequency specific spatial peak of feature importance. Moreover, we note that the impact on STN-LFP features varied significantly between these subjects. While in some cases only a few strong STN-LFP features are selected (subject S000), there were other cases in which the associated weight for the subcortical features was low (subject S008). An intermediate case is also shown in Figure 5, in which most of the STN-LFP features are chosen by the model with different weights (subject S008). Generally, the shape of the spatial patterns differs across subjects and frequency bands. Here, high-frequency oscillations have limited spatial spread (the component contributes to recordings from few electrodes, and thus abrupt changes are observed in the spatial patterns) while low-frequency oscillations present higher spatial spread, meaning that the source is contributing to recordings from several electrodes [22,23]. The individual spatial information cannot be inferred a priori, but by visual inspecting the spatial patterns, feature importance, and model decoding could be anticipated.

### 3.3 The importance of individualized subject specific decoding models

For clinical standardization and out-of-the-box use of machine learning powered brain computer interfaces in the future, the development of cross-patient decoding approaches would be beneficial. However, the results of our study highlight the individualized spatio-spectral patterns that make such an across-patient implementation difficult. Since spatial filters were used to extract the band-power features, we could investigate the neural activity at each electrode location in each frequency band, by plotting the corresponding spatial pattern (Figure 5). The spatial pattern at each frequency band reflects the mapping (in strength and sign) of the movement task-related source at each electrode. Although some brain maps across subjects share similarities, each subject has its own spatial map at each frequency band. Ultimately, for clinical brain computer interfaces, the individual spatio-spectral activity patterns will be more informative for a patient specific precision medicine approach to adaptive deep brain stimulation.

### 3.4 Limitations

Several limitations of this study are worth noting. First, we based our analyses on data obtained with a grip-force task, which describes just one type of motor behavior. Second, the data was recorded during DBS implantation surgery, when participants were without medication or neurostimulation, an otherwise unnatural state relative to typical condition. Given that such conditions change the state of recorded brain activity, further research is needed to account for these potential sources of variability. Thirdly, in general, the number of contacts in both STN and cortex recordings was sometimes too low to appreciate the spatial information of the signals. Lastly, our numerical experiments were based on a single session per patient. Considering the lack of stationarity in the electrophysiological brain data, future models ideally should be tested in several recording sessions, acquired at different days.

### 3.5 Clinical relevance

While grip-force decoding per se, may not be required for adaptive stimulation paradigm, we would like to argue that it is a very good application to investigate decoding methods, because it allows a fine-grained analysis of brain signals and behavior. Importantly, grip-force represents movement vigor, which is known to be modulated by the basal ganglia and the STN [24] and can be impaired in PD. Independent of the specific target variable of grip-force, the results shown here are likely transferrable to other domains of symptom and behavior decoding, which may be augmented by spatio-spectral methods in specific patients.

In conclusion, we have shown that movement decoding could benefit from i) the use of the spatial information and ii) multimodal brain recordings. Our results suggest that by combining two recording modalities both the decoding capacity of the model and model robustness can be improved in some cases, but an individual assessment of features and model performances is required to achieve optimal decoding performances. The high impact contribution of cortical recording for movement decoding supports the utility of ECoG for future invasive bidirectional BCI devices.

## 4 Methods and Materials

### 4.1 Data

The dataset used in this study (available upon request) corresponds to that from a previously published study [25] and is comprised of subthalamic LFP and subdural ECoG recordings simultaneously acquired from 11 PD patients (1 female, mean age ±SD = 60.1 ±8.3 years, disease duration ±SD = 9.8 ±4.0 years) undergoing DBS implantation surgery. All subjects were recommended for surgery by a multidisciplinary review board and provided written informed consent. The study was approved by the Institutional Review Board of the University of Pittsburgh (IRB Protocol #PRO13110420). UPDRS Part III scores for the off-medication conditions were collected in a time period of 1-3 months prior to surgery by movement disorder neurologists. Antiparkinsonian medications were held off for at least 12 hours prior to intraoperative testing.

Subjects were instructed to press a hand-grip force transducer with either their right or left hand after a visual cue appeared. The laterality of the movement (contra- or ipsilateral) was annotated with respect to the electrodes’ hemisphere localization. A trial was considered successful if the subject was able to maintain for at least 100 ms with the indicated hand at least 10% of their maximum voluntary grip force. Each trial was followed by a variable inter-trial interval of 500–1000 ms. During the task, subjects were fully awake. No anesthetic agents were administered for at least 1 hour prior to the task procedure. No medication was given during the task.

ECoG recording were acquired using 6, 8, 28 or 32 contacts electrode strips (Ad-Tech, Medical Instrument Corporation), placed as close as possible to the hand knob area through the burr hole used for DBS lead implantation [26]. Ground and reference electrodes were placed in the shoulder and mastoid, respectively. LFPs from the STN were recorded using a clinical four contacts DBS lead (model 3389, Medtronic). Data was sampled at 1000 Hz and band-pass filtered (0.3–250 Hz), using a Grapevine neural interface processor (Ripple Inc.). STN-LFP data was re-referenced offline to a bipolar montage by referencing each contact to its immediate neighbor, thus three new bipolar channels were generated after this procedure.

All the analogical signals, corresponding to the grip-force output, were processed offline to remove baseline drift, by estimating the baseline using the optimization problem proposed in [27] and then subtracting it from the original output. After this procedure, the grip-force output was a continuous signal with zero (rest) or positive values (movement), where a value different from zero corresponds to the grip-force applied by the subject.

### 4.2 Spatio-spectral multiple recording site decoding pipeline

A subject-specific invasive neurophysiology movement decoding model that combines a filter-bank analysis with a spatial filtering approach was implemented. Data segments of 1000ms were decomposed in eight frequency bands. Then the supervised SPoC spatial filtering approach was applied at each frequency band, extracting one spatio-spectral feature per frequency band considered. The resulting feature vector was used as input to a GLM with Poisson-like distribution sparse regression. A visual representation of the model is shown in Figure 2. The decoding pipeline consists of three online-compatible main steps: i) data pre-processing, ii) feature learning and iii) movement decoding.

#### 4.2.1 Online-compatible data pre-processing

Considering real-world applications, pre-processing steps should account for online real-time decoding challenges, that is, data is only available in packets, and future packets from the time point of decoding are not available to the decoder or processing pipeline. Thus, in this work, consecutive epochs of 1000 ms in step of 100 ms were extracted from the brain recording in order to continuously decode the grip-force target. Common average reference was applied to the extracted ECoG segments. All epochs were notch (60/120/180 Hz) and band-pass filtered in eight frequency bands of interest *θ* : [4-8] Hz, *α*: [8-12] Hz, low*β*: [13-20] Hz, high*β*: [20-35] Hz, all*β* [13-35] Hz, low*γ*: [60-80] Hz, high*γ*: [90-200] Hz, and all*γ*: [60-200] Hz). The pre-processed brain recordings were arranged in a four-dimensional array accounting for the number of extracted epochs *N_t_*, the number of channels *N_c_*, the numbers of sample point per epoch *N_s_* and the number of filter-bands *N_f_*. In the case of the target variable, in accordance with the 1000 ms time window length extracted in steps of 100 ms, it was downsampled by selecting the 100^*th*^ sample point from the processed grip-force. These online-compatible pre-processing steps were made by the py_neuromodulation package (https://github.com/neuromodulation/py_neuromodulation).

#### 4.2.2 Feature extraction via source power comodulation

Electrophysiological recordings can be modeled as a linear and instantaneous superposition of neural sources [28,29]. Let 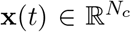 be the brain recordings in the sensor space (raw data) at time *t*, where *N_c_* denotes for the number of channels. Let 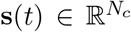 be the sources (or components) and let 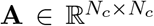 be the mixing matrix, whose *i^th^* column vector **a**_*i*_ is what is known as *spatial pattern*. Considering additive noise, the following definition holds to represent the generative model:

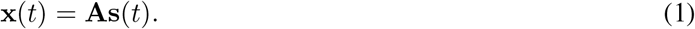

In the context of spatial filtering, the objective is to estimate **s**(*t*) and thus, transform the signal from the *sensor* to the *source* space. This sensor-to-source transformation can be obtained by means of the so-called blind source separation methods, as follows:

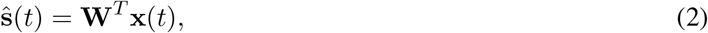

where **W** = [**w**_1_,…, **w**_*i*_,…, **w**_N_c__] is a *N_c_* × *N_c_* demixing matrix whose *i^th^* column vector corresponds to what is known in the literature as *spatial filter*. Each of the spatial filter is meant to extract the signal from one source while suppressing the activity of the others, such that the resulting projected signal is a close approximation of the original source signal [14]. In the particular case of the SPoC algorithm, method simultaneously discovered by [14,30], the information contained in the target variable is used to guide the decomposition. Denoting **z**(*t*) to the target variable (grip-force movement), the first SPoC filter is found by:

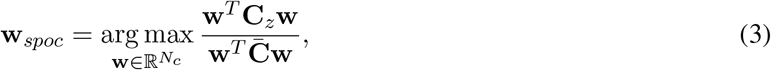

where 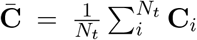 is the Euclidean average covariance matrix across the *N_t_* data segments and 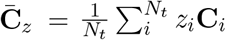 is the weighted average covariance matrix. The rest of the filters are obtained by solving a generalized eigenvalue decomposition problem [14]. As shown in [13], the matrix **W**_*spoc*_ recovers the inverse of mixing matrix **A** defined in (1).

When data has been band-pass filtered, the power of the projected signal **w**^*T*^**x**(*t*) approximates the target function **z**. Thus, after applying SPoC spatio-spectral features can be extracted by

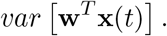

In this work, the SPoC method was applied at each frequency band considered. Each band-power feature was extracted from the projected signal in the first SPoC component, taking the entirely 1000 ms for the *θ* band, last 500 ms for *α*, last 330 ms for the *β* bands and last 100 ms for the *γ* bands. Thus, at the end of the procedure eight (8) spatio-spectral features were computed. It is timely to mention here that while offline learning of the spatial filters **W**_*spoc*_ is needed, once such a matrix is learned, the spatio-spectral features can be extracted in real-time by projecting the signals through **W**_*spoc*_.

#### 4.2.3 Generalized regularized linear models

Let 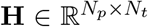 be the matrix of *N_p_* predictors (features), where *N_t_* is the number of extracted epochs. Let 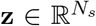 be as before, the target variable. Traditional linear model assumes that the distribution of the output is normal 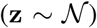 and thus, it can be modeled as a linear combination of the predictors and suitable weights, that is:

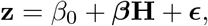

where the model parameters *β*_0_, 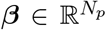 can be estimated using ordinary least squares or its regularized versions. In the particular case of the elastic-net penalty (enet), a compromise between the *ℓ*_1_ and *ℓ*_2_ norm of the solution is imposed, and thus the solution vector and intercept can be found by solving the following unconstrained regularized problem:

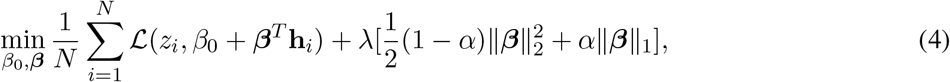

where 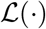 is the loss function aimed to be minimized, **h**_*i*_ is the *i^th^* column vector of **H**, and λ and *α* are positive constants called regularization parameters. In particular, α ∈ [0,1] and thus it balances between the Ridge regression (*α* = 0) and LASSO regression (*α* = 1).

When working with non-normally distributed outputs, we can extend this approach by means of generalized linear models (GLMs). The Poisson distribution is a discrete probability distribution that expresses the probability of a given number of events occurring in a fixed interval of time. Since the grip-force target is a non-negative output which accounts for rest and movement events, we assume here, from practical reasons, that **z** can be modeled as coming from a Poisson-like distribution. Under this assumption, a log - `exp` relationship between the predictors and the output exists. Using the softplus formulating proposed by [31], the loss function 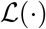 in (4) is given by:

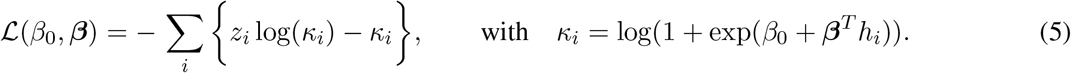

### 4.3 M_odel training and evaluation_

In order to investigate the practical use of the spatial filters for the development of aDBS, different computational experiments were conducted. Since the dataset comprised bilateral movements (contra- and ipsilateral to the electrodes’ position on the respective hemisphere), one decoding model was built for each subject at each movement laterality. A 5-fold non-shuffled cross-validation procedure was performed, where the coefficient of determination *R*^2^ was used for measuring the performance of the predicted grip-force when compared to the true grip-force output. The SPoC algorithm was applied as a feature extraction method at each frequency band considered. The demixing matrix *W_spoc_* was learned using the training set in each cross-validation fold, and features from the unseen testing set were extracted using these estimated filters. One spatial filter was used to compute the feature at each frequency band, leading to eight spatio-spectral features at the end of the filter-bank analysis. Features were concatenated and z-score scaled. In order to avoid any circularity, the statistics (mean and standard deviation) were estimated in the training set and then applied to scale the unseen testing test. These features corresponded to the inputs to the Poisson-like GLM with enet penalty regression model. In order to equally balance the Ridge and Lasso penalty, we set the *α* parameter of (4) equal to 0.5. The regularization parameter λ was searched via Bayesian Optimization [32] over a 3-fold non-shuffled cross-validation on training data. Statistical analyses were performed via the non-parametric Friedman test and the post-hoc Nemenyi test at level of significance 0.05 [33].

#### 4.3.1 Computing environment

The corresponding Python source codes developed for this work are publicly available at https://github.com/Brain-Modulation-Lab/Paper_SpatialPatternsMovementDecoding. We used the py_neuromodulation package (https://github.com/neuromodulation/py_neuromodulation) for implementing the online-compatible pre-processing steps, the MNE-Python library [34](https://mne.tools) for implementing SPoC, the pyglmnet package (https://pypi.org/project/pyglmnet/) for running the GLM [35], scikit-learn for constructing pipelines [36](https://scikit-learn.org), and the Bayesian Optimization package [37](https://github.com/fmfn/BayesianOptimization) for finding the optimal regularization parameter.

## 5 Acknowledgments

We would like to thank the subjects for their generous gift of time in the operating room and additional experimenters who acquired and organized the data. In addition, we would like to thank Denis A. Engemann for suggesting the use of a generalized linear model with non-negative distributions. The present manuscript was supported through a US-German Collaborative Research in Computational Neuroscience (CRCNS) grant, funded by the Federal Ministry for Research and Education (Germany) and the National Institutes of Health (United States, R01NS110424). In addition this manuscript was partially funded by the Deutsche Forschungsgemeinschaft (DFG, German Research Foundation) – Project-ID 424778381 – TRR 295.

## Additional Information

### Author contribution

**Victoria Peterson:**Conceptualization, Methodology, Software, Validation, Formal analysis, Writing - Original Draft, Visualization. **Timon Merk**: Software, Data Curation, Writing - Review & Editing. **Alan Bush:** Conceptualization, Investigation, Data Curation, Writing - Review & Editing. **Andrea A. Kühn:**Conceptualization, Writing - Review & Editing, Project administration, Funding acquisition. **Vadim Nikulin:** Conceptualization, Methodology, Writing - Review & Editing. **Wolf-Julian Neumann:** Conceptualization, Writing - Review & Editing, Supervision, Project administration, Funding acquisition. **Mark Richardson:** Conceptualization, Investigation, Writing - Review & Editing, Supervision, Project administration, Funding acquisition.

